# Circadian transcriptome in cultured cells depends on synchronisation method and differs from *in vivo* condition

**DOI:** 10.64898/2026.07.20.739603

**Authors:** Michal Dudek, Cátia F. Gonçalves, Judith A. Hoyland, Qing-Jun Meng

**Affiliations:** Manchester Cell Matrix Centre, Faculty of Biology, Medicine and Health, University of Manchester, M13 9PT, UK; Centre for Biological Timing, Faculty of Biology, Medicine and Health, University of Manchester, M13 9PT, UK; Division of Cell Matrix Biology and Regenerative Medicine, Faculty of Biology, Medicine and Health, University of Manchester, Manchester Academic Health Science Centre, Manchester, M13 9PT, UK; NIHR Manchester Biomedical Research Centre, Central Manchester Foundation Trust, Manchester Academic Health Science Centre, Manchester, M13 9PT, UK

## Abstract

*In vitro* synchronisation is widely used to study circadian clocks in cells, but whether cultured cells recapitulate tissue-level rhythmic outputs remains unclear. Articular cartilage provides a useful model to address this because chondrocytes are the only resident cell type. Here, we compared circadian time series transcriptomes between primary mouse chondrocytes synchronised by heat shock, dexamethasone, or osmotic stress and *in vivo* cartilage tissue. All three stimuli robustly synchronised core clock gene rhythms but produced distinct circadian phases and markedly different rhythmic transcriptomes, depending on the synchronizer. Heat shock, dexamethasone, and osmotic stress yielded 5255, 2008, and 879 transcripts classified as rhythmic, respectively, in primary chondrocytes, yet only 64 genes were shared across the three *in vitro* datasets, and only 15 were shared when *in vivo* cartilage transcriptome was included. Pairwise comparisons between synchronizers revealed some statistically enriched overlaps, but only marginally above chance, and shared genes showed limited conservation of circadian phase. Functional enrichment analysis also revealed stimulus-dependent rhythmic programmes with modest pathway-level overlap. These findings indicate that circadian output in cultured cells is shaped by the synchronising cue and cellular microenvironment. We also present BodyClocks.org, an interactive resource implementing this comparative framework across a curated collection of circadian transcriptomic datasets.

## INTRODUCTION

The adaptation of life to the Earth’s 24-hour axial rotation has resulted in the evolution of the circadian timing system, a highly conserved, hierarchical network allowing organisms on Earth to anticipate and align physiological functions with daily environmental changes [1]. In mammals, the central circadian clock in the suprachiasmatic nucleus synchronises a network of peripheral clocks present in most tissues and cell types [2]. At the cellular level, these clocks are generated by conserved transcriptional-translational feedback loops in which transcription factors CLOCK and BMAL1 activate, while PER and CRY repress, rhythmic gene expression. Although this core molecular clock machinery is conserved between cell types, the downstream clock-controlled genes (CCGs) are highly tissue-specific [3]. Depending on the tissue, the physiological state of the organism, and the stringency of the bioinformatics algorithms employed, CCGs can represent anywhere from 3% to 20% of the actively transcribed genome of a given tissue. It is through these CCGs that the circadian clock temporally segregates incompatible metabolic pathways and coordinates tissue-specific functions to the diurnal cycles [4].

Articular cartilage is a specialised load-bearing connective tissue, providing a smooth and lubricated surface for joint articulation and load transmission. Its resident chondrocytes, the only cell type in the tissue, are embedded in a dense extracellular matrix and are solely responsible for maintaining the balance between matrix synthesis and degradation required for lifelong tissue integrity [5]. We and others have demonstrated that chondrocytes possess a functional, cell-autonomous circadian clock [6–8]. Real-time bioluminescence recordings from *PER2::LUC* transgenic reporter mice demonstrate that cartilage explants exhibit self-sustained, robust circadian oscillations *ex vivo*. Furthermore, *in vivo* transcriptomic profiling has revealed that approximately 3.9% of the cartilage genome is rhythmically transcribed, featuring genes critical for tissue homeostasis, cellular survival, and TGF-β signalling [7]. Disruption of the chondrocyte clock, including deletion of *Bmal1*, accelerates cartilage ageing and degeneration [7]. Because peripheral clocks are semi-autonomous and susceptible to intrinsic molecular noise, their TTFLs will require daily entrainment by *zeitgebers* (time cues) to maintain phase coherence at the tissue level [9]. In this regard, the anatomical isolation of articular cartilage poses a unique entrainment challenge. Lacking direct innervation and vascular supply, chondrocytes are insulated from rapid systemic neural signals. Instead, they must discern the time of day by sensing diffusible factors (such as hormones) and, crucially, the local physicochemical fluctuations of their unique biomechanical niche. So far, three categories of *zeitgebers* have been identified as synchronisers of the chondrocyte clocks *in vivo*: endocrine, thermal, and mechanical/osmotic signals [10–12].

A breakthrough in the chronobiology field was the discovery that even cultured cells, such as fibroblasts, contain functional circadian clocks which can be studied in culture for many days [13]. This cellular approach has paved the way for biochemical and genetic dissection of the circadian clock mechanisms, providing significant insights into the molecular intricacies of the clockwork at cellular level, and enabled the search for new clock components, modifiers and drugs [14–16]. A common laboratory approach for studying peripheral clocks is to culture established cell lines or primary cells and expose them to an acute synchronising stimulus before time-course sampling [17–19]. This reductionist strategy has been essential for defining core clock mechanisms and input pathways, but it also removes cells from their native physiological environment. Culture conditions have been shown to substantially alter the transcriptional identity of primary cells [20,21]. Therefore, question remains as to whether synchronised cells *in vitro* recapitulate the rhythmic clock output programmes present in intact tissues. Therefore, using this cellular approach as a proxy to infer rhythmic outputs of a particular cell type *in vivo* is potentially problematic, because this approach relies on the idea that once the core TTFL is synchronised by any valid *zeitgeber*, the clock will faithfully drive its innate, pre-programmed suite of “tissue-specific” clock-controlled genes. This assumes a linear, unidirectional relationship where the synchronisation method merely starts the pendulum swinging, and the downstream gearwork remains constant and passively responds. However, emerging evidence challenges this static view. The core clock proteins do not function in isolation. CLOCK and BMAL1 frequently cooperate with lineage-specific or stimulus-activated co-transcription factors, and their binding is dependent upon the chromatin accessibility landscape of the cell [22–24]. Chondrocytes are indeed highly responsive to their microenvironment. The transition from a 3D mechanically active matrix *in vivo* to a static, 2D rigid substrate induces rapid dedifferentiation and fundamentally alters basal gene expression [25].

However, the complex and diverse cellular composition (including blood vessels and nerves) of most tissues makes it difficult to determine whether rhythmic outputs observed *in vitro* faithfully reflect cell-type-specific rhythms *in vivo*. In this regard, articular cartilage provides a uniquely simple model because unlike many other tissues, chondrocytes are the only resident cell type in the avascular and aneural cartilage tissue. On the other hand, these chondrocytes are exposed *in vivo* to endocrine, thermal, mechanical, and osmotic time cues, which can be modelled in cell culture. Here, we used primary mouse chondrocytes and *in vivo* cartilage transcriptomic datasets to test whether different synchronising stimuli produce a common chondrocyte circadian output or distinct stimulus-dependent rhythmic programmes. We compared rhythmic gene identity, phase relationships, and pathway enrichment across chondrocytes synchronised *in vitro* by heat shock, dexamethasone, or osmotic stress, and against the rhythmic transcriptome of cartilage *in vivo*. To facilitate interpretation of these comparisons and extend the same analytical framework to public datasets in many other tissues of mouse and baboon, we developed BodyClocks, a freely available, open-source web resource for exploring circadian transcriptomes at gene, pathway, network, and dataset-comparison levels. We envisage this resource to be of broad interests for comparative studies of circadian regulated pathways, programs and networks in different tissues and species.

## METHODS

### Animal husbandry

All experimental procedures were carried out in accordance with the Animals (Scientific Procedures) Act of 1986 under Home Office license PP7202624 and with approval of local ethics committee. Mice were group housed under a 12 hours light/12 hours dark regimen (lights on at 7 AM and off at 7 PM) with ad libitum access to standard rodent chow and water. All animals were bred by the Biological Services Facility at the University of Manchester and euthanised by CO_2_ and cervical dislocation between zeitgeber time 5.5 (ZT5.5) and ZT6.5 (between 12.30 PM and 1.30 PM). PER2::LUC mice of both sexes carrying the firefly luciferase gene fused in frame with the 3’ end of Per2 were utilised in all studies [26].

### Primary articular chondrocytes

Primary articular chondrocytes were isolated from five-day-old mice as previously described [27] with minor modifications. In brief, articular cartilage was isolated from femoral heads, femoral condyles and tibial plateaus of mice from one litter. Explants were rinsed with 1 × phosphate buffered saline (PBS, SigmaAldrich) and incubated in digestion solution (3 mg mL-1 collagenase D [Roche] in DMEM/F-12 31330038 [Gibco]) for 45 min at 37 °C with intermittent vortex mixing to dislodge soft tissues. Subsequently, samples were diced using a sterile scalpel and incubated overnight at 37 °C in the digestion solution. Cell aggregates were dispersed by successively passing the solution through 25- , 10- and 5-ml Pasteur pipettes. The cell suspension was filtered through a 70 µm cell strainer (EASYstrainer, Greiner Bio-One) and then centrifuged for 10 min at 400 ×g. Finally, the pellet was resuspended in DMEM/F-12 supplemented with 10% FBS (Gibco), and 100 units penicillin/100 µg mL-1 streptomycin (Sigma-Aldrich). Primary articular chondrocytes were used up to passage 1.

### Treatment conditions

For dexamethasone and heat shock experiments, all cultures were initially synchronised with dexamethasone to establish comparable baseline clock oscillations before secondary clock resetting. This step was necessary as we found that heat shock synchronisation resulted in the highest amplitude when applied at projected peak of PER2 as described in [11]. A “washout” period of 112 hours was introduced to allow acute dexamethasone-responsive transcriptional effects to subside and PER2::LUC oscillations were barely detectable before the application of heat-shock or second dexamethasone resetting stimulus and subsequent time series sampling for RNA-seq. Specifically, to synchronise circadian rhythms, chondrocytes at 90% confluence in 35 mm cell culture dishes were incubated with 100 nM of dexamethasone for 60 min. Afterwards, media was changed and dishes were sealed with 40 mm glass cover slips (Thermo Fisher Scientific) and high vacuum grease (Dow Corning). Sealed dishes were placed in the Lumicycle bioluminescence recorder (Actimetrics) or an incubator (for mRNA time-course) at 37 °C for 112 hours. At 112 hours half the dishes (heat shock group) were moved to an incubator at 43 °C for 60 min. The other dishes (dexamethasone group) remained in an incubator at 37 °C and a second pulse of 100 nM dexamethasone were added to the culture media approximately 15 min prior to the end of the heat pulse in the other group. All dishes were then returned to the Lumicycle (for recording) or incubator at 37 °C and the first time-course sample was collected four hours later, after which chondrocytes were harvested for RNA isolation every four hours for 48 hours. This experiment was repeated three times in chondrocytes isolated from distinct litters. Synchronisation by osmotic stress is described in [12] study from which the circadian dataset was used here for comparison (E-MTAB-11040). The cartilage circadian dataset (E-MTAB-3428) was first published in [7].

### Mouse articular chondrocytes library construction and sequencing

Total RNA was isolated using the PureLink RNA Mini Kit (Thermo Fisher) as per manufacturer’s instructions. 50 ng of RNA per sample was used for library generation. mRNA library preparation and sequencing were performed at the University of Manchester’s Genomic Technologies Core Facility. Libraries were generated using the TruSeq Stranded mRNA assay (Illumina) according to the manufacturer’s protocol. Libraries were sequenced (2 × 75 bp paired-end reads) in the HiSeq 4000 instrument (Illumina). The output data was demultiplexed (allowing one mismatch) and BCL-to-Fastq conversion performed using Illumina’s bcl2fastq software v2.20.0.422.

### Read pre-processing, filtering and mapping

Quality control and pre-processing of FASTQ files was performed using FastQC v 0.11.3 and MultiQC v1.8. Sequence adapters were removed and reads were quality trimmed using Trimmomatic v0.39 [28]. Reads were then mapped to the Ensembl GRCm38/mm10 mouse genome and counts per gene were calculated using STAR v2.7.3a with annotation from GENCODE M25 [29].

### Bioinformatic analysis

#### RNA sequencing data processing and circadian rhythm analysis

Raw RNA sequencing count data were processed using DESeq2 (v1.50.2) [30]. Prior to analysis, sample quality was assessed with Principle component analysis (PCA) and outlier samples were excluded (heat shock timepoint 24h sample b, and dexamethasone timepoint 28h sample b). Batch correction was performed using ComBat-seq [31] from the sva package prior to DESeq2 normalisation of the heat shock and dexamethasone datasets. The batch variable was defined as the experimental time-course replicate, corresponding to the three independently generated 48-hour time courses. To preserve circadian time-dependent expression patterns during batch correction, the ComBat-seq group variable was defined by sampling time point, so that samples collected at the same circadian time across the three experimental batches were treated as the same biological group. This design allowed batch-associated variation to be adjusted while retaining the rhythmic temporal structure required for downstream RAIN analysis. A minimum mean expression threshold of 50 raw counts was applied to all datasets to filter lowly expressed genes. Rhythmically expressed genes were identified using the RAIN (Rhythmicity Analysis Incorporating Non-parametric methods) algorithm [32], applied to expression data across a 48-hour time-course sampled at 4-hour intervals with a period of 24 hours, using the longitudinal and independent methods for chondrocyte and cartilage datasets respectively, with peak borders set to [0.3, 0.7]. Raw p-values were corrected for multiple testing using the Benjamini-Hochberg procedure, and genes with an adjusted p-value (BH.Q) < 0.05 were classified as significantly rhythmic. Gene identifiers were annotated with MGI symbols, gene names, and functional descriptions by querying the Ensembl BioMart database (*Mus musculus*) via the biomaRt R package, preferentially using MGI symbols with fallback to external gene names where MGI symbols were unavailable. Results were visualised as expression heatmaps ordered by peak phase using the ComplexHeatmap package.

#### Functional enrichment and protein-protein interaction network analysis

For functional characterisation of rhythmically expressed genes, protein-protein interaction networks and functional enrichment analyses were performed using the STRING database [33] (v12.0, *Mus musculus*, confidence score ≥ 700) via the STRINGdb R package. Enrichment was assessed across four functional annotation categories: KEGG pathways, Gene Ontology Biological Process terms, Reactome pathways, and WikiPathways, with results filtered at a false discovery rate ≤ 0.05. Ensembl gene identifiers were used as primary query inputs, as these yielded more complete mappings than MGI symbols. Gene identifier mapping between Ensembl IDs, STRING preferred names, and MGI symbols was performed using a custom dictionary built from BioMart [34] and STRING mapping outputs. Processed results, including interaction networks, enrichment annotations, and per-gene phase information, were saved as RDS files for use in the BodyClocks platform.

#### Comparative overlap analysis between datasets

To compare the rhythmically expressed transcriptomes across experimental conditions, pairwise overlaps between the osmotic stress, dexamethasone, heat shock, and cartilage circadian time course datasets were assessed. Rhythmically expressed genes were identified in each dataset by filtering for BH.Q < 0.05, with a secondary analysis performed at the more stringent threshold of BH.Q < 0.01. The statistical significance of pairwise rhythmic-gene overlaps was evaluated using a one-sided hypergeometric test, assessing whether the observed number of genes rhythmic in both datasets exceeded that expected by chance. For each pairwise comparison, the background universe was defined as all genes that passed expression filtering and were tested for rhythmicity in both datasets, i.e. the intersection of the two datasets’ expressed/tested gene sets. A one-sided test was used because the prespecified hypothesis was enrichment of shared rhythmic genes above chance expectation, rather than either enrichment or depletion.

For each dataset pair, genes rhythmic in both datasets (BH-adjusted p < 0.05) were compared by their clock-relative phase (CRP), defined as each gene’s circadian phase expressed relative to *Bmal1* within that dataset. Two genes were considered phase-concordant if their circular distance, the minimum arc on a 24 h clock, was ≤ 4 h (one timepoint resolution). To test whether the observed proportion of concordant genes exceeded chance, a permutation null distribution was generated by randomly shuffling the CRP values of one dataset across genes (1,000 permutations) while holding the other fixed and recomputing the concordant proportion each time. For pairwise comparisons, the theoretical expectation for a randomly placed phase to fall within a ±4 h window on a 24 h circle is 8/24 = 33.3%. However, because observed rhythmic genes are not uniformly distributed across phase, significance was assessed using a permutation null. For the three-way comparison, the expected concordance is lower because all three pairwise distances must fall within the threshold; therefore, a separate permutation null was generated. The empirical *p*-value was the proportion of permutations yielding a concordant fraction ≥ the observed value, and Benjamini–Hochberg correction was applied across the six pairwise comparisons. For the three-way analysis (genes rhythmic in osmotic stress, dexamethasone, and heat shock treatments simultaneously), a gene was classified as concordant if all three pairwise circular distances were ≤ 4 h. Osmo CRP values were held fixed while Dex and HS values were permuted independently (1,000 permutations), and the *p*-value was calculated as above.

Functional enrichment of rhythmic genes (RAIN BH-adjusted *p* < 0.05) was performed locally using mouse KEGG, Reactome and WikiPathways gene - term annotations. Pathways containing at least five background genes were tested for over-representation using one-sided hypergeometric tests, with BH correction within each dataset and annotation category. For each pairwise comparison, the primary enrichment background was the intersection of all genes tested for rhythmicity in both datasets, restricted to genes mapped to STRING. Similarity between significant term sets (FDR ≤ 0.05) was quantified using the Jaccard index. Significance was assessed using 1,000 permutations in which independent gene sets matching the observed rhythmic-gene set sizes were sampled from the corresponding background. Because random gene sets rarely produced FDR-significant pathways, terms were ranked by nominal enrichment *P*-value and the top *N* terms retained, where *N* matched the observed significant-term list size. Empirical *P*-values were calculated as (*b* + 1)/(*B* + 1) and BH-adjusted across the 18 dataset-pair–annotation-category comparisons. The analysis was repeated using all STRING-mapped mouse genes as a genome-wide background, corresponding more closely to default STRINGdb enrichment. An exact term-level hypergeometric test was included as a sensitivity analysis.

Complete analysis pipeline including figures, tables and data files for the BodyClocks application is available at GitHub (https://github.com/Michal0110/BodyClocks_data).

#### Web Application Design

The BodyClocks web application was developed using R Shiny (version ≥1.11.1) as an interactive platform for exploration of circadian RNA sequencing datasets. The application follows a modular architecture, with a reusable explorerServer/explorerUI module pair handling the Circadian Explorer tab and allowing future expansion with other specialised tabs. Protein–protein interaction networks were pre-computed using the STRINGdb database and rendered interactively using the visNetwork package [35], with nodes representing genes and edges weighted by interaction confidence. Functional enrichment annotations, including GO Biological Process, KEGG Pathway, Reactome, and WikiPathways terms, were pre-computed and linked to network clusters, enabling users to highlight gene sets of interest directly within the network. Gene expression data are displayed as temporal profiles fitted with nonlinear least-squares (NLS) sine and cosine models using a multi-start strategy to identify the best-fitting oscillatory curve. Model selection was based on a composite score combining residual deviance and deviation from the expected circadian frequency. The data table, built with the DT package, supports multi-row selection for gene expression plotting, and all visualisations are rendered dynamically using ggplot2. The methods described in the above paragraph “*Comparative overlap analysis between datasets*” are implemented as a separate interactive tab that allows pairwise comparisons between any two datasets of the same species. The application is deployed as a standard Shiny application at www.BodyClocks.org and the source code and Docker container for local deployment are available at GitHub (https://github.com/Michal0110/BodyClocks_app).

## RESULTS

### Different synchronising cues generate robust but distinct rhythmic outputs in primary chondrocytes

We have previously established that osmotic stress [12] and heat shock [11] are potent clock resetting stimuli for cartilage *ex vivo*, while dexamethasone remains one of the most widely used synchronisation methods in circadian studies [36]. To systematically evaluate the impact of synchronisation modality on the circadian output of a peripheral clock, we conducted an RNA-sequencing time-series investigation using mouse primary articular chondrocytes. We first tested whether the three synchronising cues produced comparable circadian transcriptomes in primary chondrocytes. All three methods resulted in re-synchronisation and robust circadian rhythms of PER2::LUC clock reporter in chondrocytes (Figure 1A). RNA sequencing of 48-hour circadian time series revealed that all three methods induced transcriptome-wide rhythms. Osmotic stress, heat shock and dexamethasone induced 879, 5255 and 2008 rhythmic transcripts, respectively (Figure 1B). The overlap between lists of rhythmic genes in the three chondrocyte datasets was only 64 genes (Supplementary Table 1) and when cartilage tissue rhythmic genes were added to the comparison, only 15 genes were common between all datasets. This common set contained largely core circadian clock genes (e.g., *Bmal1, Npas2, Cry1, Nr1d1, Per1, Per2, Per3, DBP*, and *Tef*) and their temporal pattern differed between the synchronisation modalities (Figure 1C).

**Figure 1.**
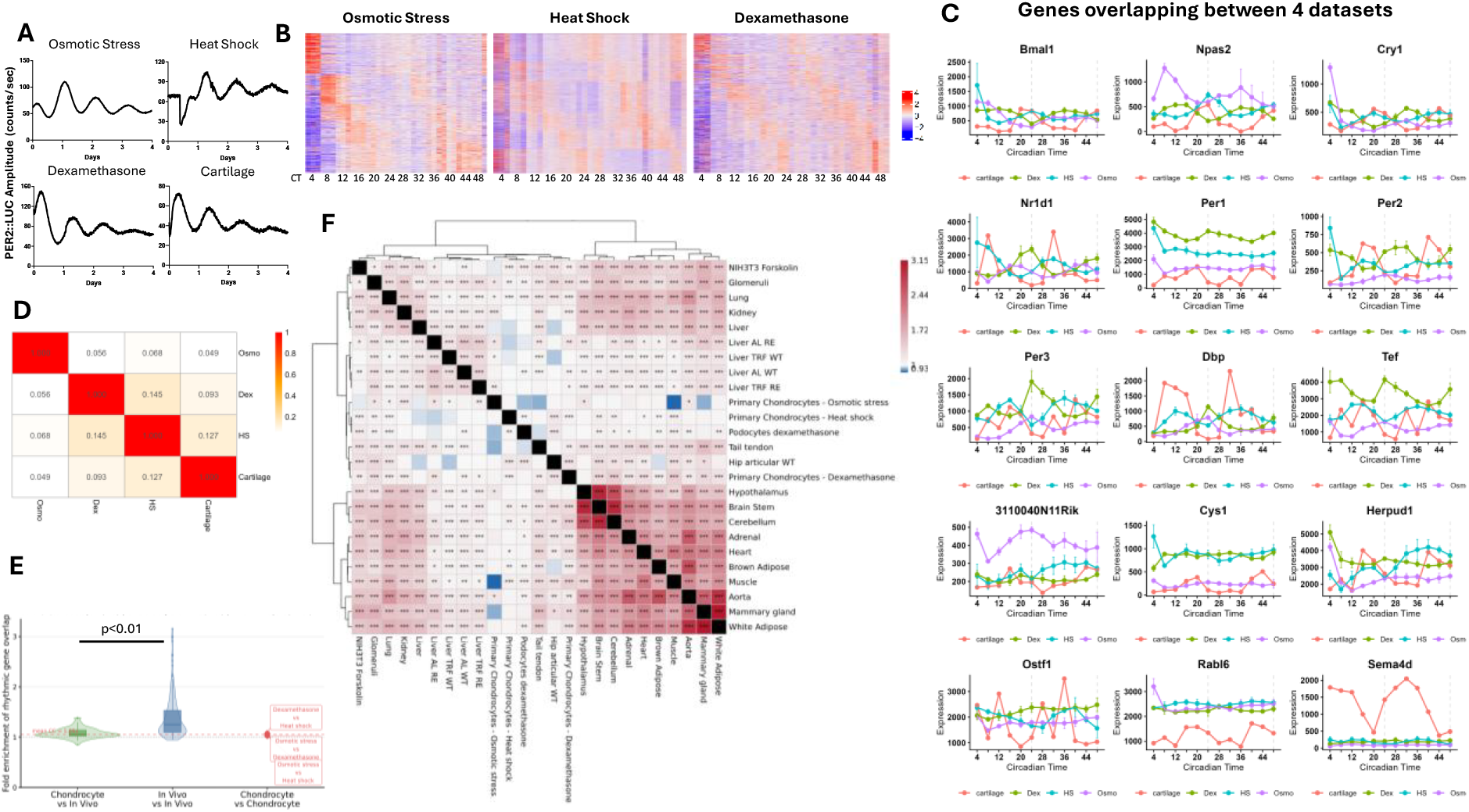
Circadian clock synchronisation with stress and glucocorticoids results in diverse rhythmic transcriptomes in mouse primary chondrocytes. **A**. Live bioluminescence recording of PER2::LUC mouse primary chondrocytes synchronised with Osmotic Stress, Heat Shock, Dexamethasone, as well as Cartilage explant with endogenous rhythm. **B**. Heatmaps of the rhythmic transcriptomes resulting from the three synchronisation methods. **C**. Gene expression patterns of rhythmic genes shared among the 4 datasets. **D**. Jaccard index illustrating similarity of rhythmic transcriptomes in pairwise comparisons between different conditions. **E**. Mean (+/- SD) fold enrichment of chondrocyte vs chondrocyte (mean = 1.06, n = 3), chondrocyte vs *in vivo* (mean = 1.08, n = 66) and *in vivo* vs *in vivo* (mean = 1.39, n =231) comparisons. **F**. Heat map of pairwise rhythmic gene overlap fold enrichment of 25 circadian mouse datasets available in the BodyClocks analysis and visualisation platform. ALT TEXT: Graphs and data showing comparisons between rhythmic gene sets induced by osmotic stress, heat shock and dexamethasone. Figures labelled A to F.

### Rhythmic genes show low overlap between synchronisation methods

We next tested whether pairwise overlaps between rhythmic gene lists exceeded those expected by chance. The analysis showed that there were more genes than there would be purely by chance in heat shock vs dexamethasone, heat shock vs cartilage and dexamethasone vs cartilage comparisons, but none of the comparisons involving osmotic stress reached statistical significance. However, the fold enrichment over random overlap even in the statistically significant comparisons (BH-adjusted p ≤ 0.05) ranged only from 1.08 fold for heat shock vs dexamethasone to 1.2 fold for dexamethasone vs cartilage (Supplementary Table 2). This corresponds to overlap similarity analysis (measured by Jaccard index where 0 is no overlap and 1 is complete overlap) of 0.145 and 0.093 for heat shock vs dexamethasone and dexamethasone vs cartilage comparisons, respectively (Figure 1D). Increasing the stringency to BH.Q < 0.01 for detection of rhythmic genes resulted in modest increase in fold enrichment. However, the overlaps were not statistically significant except dexamethasone vs heat shock (1.27 fold enrichment) and heat shock vs cartilage (1.63 fold enrichment) (Supplementary Table 2). Thus, although several overlaps were statistically enriched, the magnitude of overlap was small, indicating low similarity between rhythmic transcriptomes. Furthermore, the chondrocyte rhythmic gene sets induced by different synchronizers showed lower overlap between each other than the average overlap between any two mouse tissues. Fold enrichment was lower for chondrocyte - in vivo comparisons than for in vivo - in vivo comparisons (mean 1.083 versus 1.391; mean difference ™0.309). An exact two-sided dataset-label permutation test gave a nominal p = 0.0091. However, this did not remain significant after Benjamini– Hochberg correction across the six comparisons performed BH-adjusted p = 0.0548. (Figure 1E). When clustered based on fold enrichment against other publicly available mouse datasets, the chondrocytes form a pocket of low enrichment joined by another *in vitro* dataset of podocyte cells synchronised with dexamethasone (Figure 1F).

### Shared rhythmic genes show limited conservation of clock-relative phase

Lack of alignment of gene expression patterns in Figure 1C suggested that each synchronisation method results in a different phase of the clock oscillations. This may reflect the distinct clock resetting mechanisms by different stimuli, leading to different clock phase. Indeed, all four datasets exhibit different phase distribution of rhythmic genes (Figure 2A). Moreover, when we compared phases of core clock genes, there was little agreement between datasets (Supplementary Table 3). Core clock genes are known to show a well-preserved phase relationship among each other [37]. To account for different absolute clock phases between synchronisation methods, phases were recalculated relative to the core clock factor *Bmal1* and compared on a circular 24-hour scale. To assess whether clock-relative phase (CRP) consistency between dataset pairs exceeded chance, we computed the proportion of pairwise-overlap circadian genes maintaining a CRP distance ≤ 4 h between datasets and compared this to an empirical null distribution generated by permuting CRP values across genes (n = 1,000 permutations). Pairs involving dexamethasone, heat shock, and cartilage showed statistically significant enrichment above the permutation null of similar gene phases (Figure 2C). In contrast, all three pairs involving osmotic stress failed to exceed the permutation null indicating that osmotic stress synchronisation *in vitro* produces CRP distribution that is statistically least stable as compared to the other methods (Figure 2C). Even when the gene set was limited to the 64 genes common between the three chondrocyte datasets (Figure 2D) (presumably a highly conserved list of clock controlled genes), only 23 genes (35.9% vs. permutation null 19.4%, perm_p ≤ 0.001) oscillated with similar CRP (Figure 2E and Supplementary Table 1). Taken together, these results indicate that genes common between dexamethasone, heat shock, and cartilage pairs oscillate in similar clock relative phases, however the proportion of such genes is not overwhelming.

**Figure 2.**
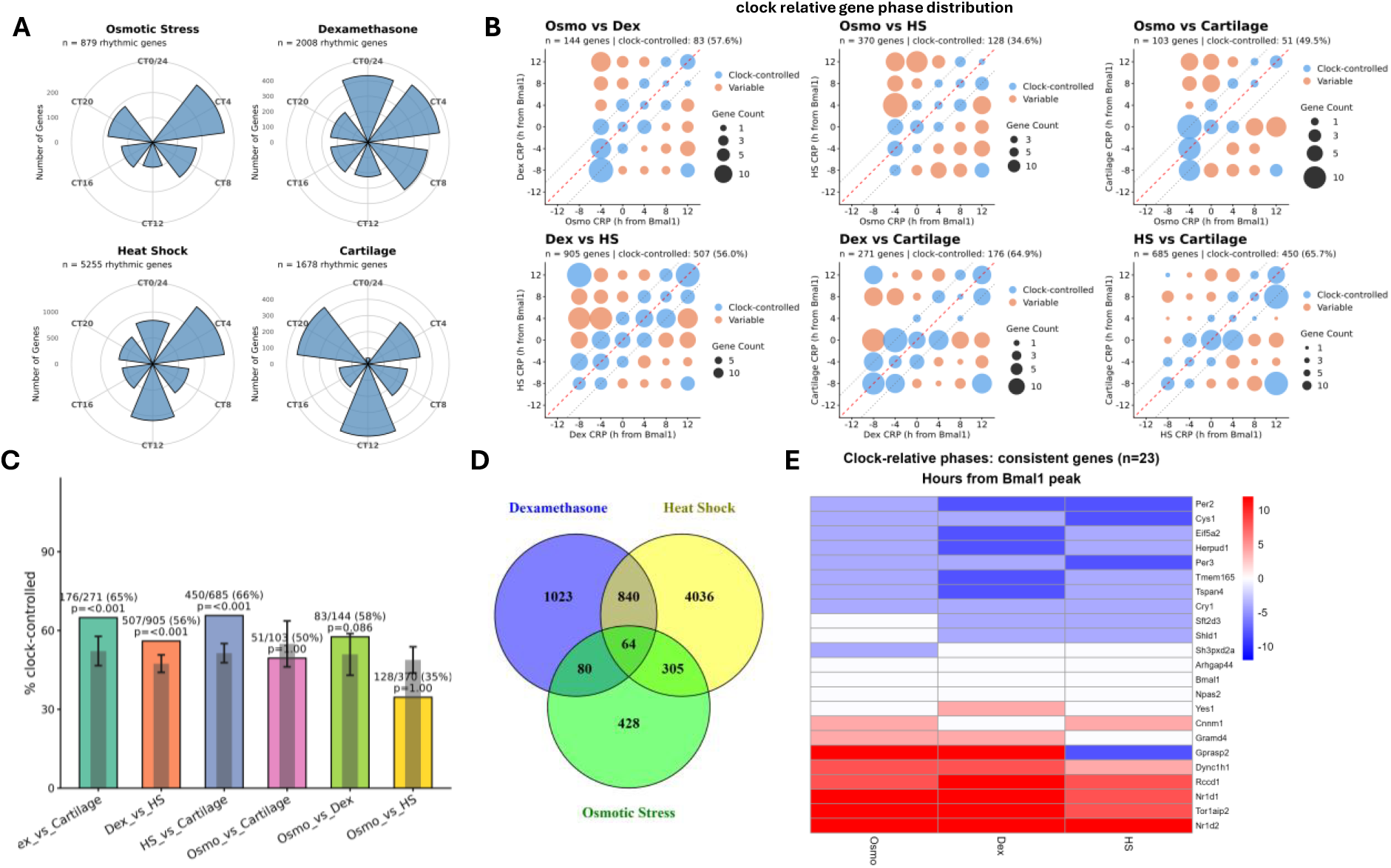
Shared rhythmic genes show limited conservation of clock-relative phase. **A**. Rose plots showing absolute phase distribution of rhythmic genes in the datasets. **B**. Scatter plots showing clock relative phase agreement (within +/- 4 h window) of rhythmic genes common between the comparison. **C**. Enrichment of genes with coherent clock relative phases over permutation null (grey bar, mean +/- SD). D. Venn diagram showing number of overlapping genes between dexamethasone, heat shock and osmotic stress datasets. **E**. Heatmap of 23 genes with coherent clock relative phase shared between the 3 conditions. Colour scale represents peak times (phase) in hours relative to *Bmal1* peak. ALT TEXT: Graphs and data showing comparison between circadian phases of genes induced by osmotic stress, heat shock and dexamethasone and rhythmic gene phases in cartilage. Figures labelled A to E.

### Pathway-level overlap remains modest and annotation-category dependent

Although the overlap of rhythmic genes between the datasets was low, we hypothesised that perhaps the rhythmic genes from different stimulus could belong to similar rhythmic pathways. To test whether there is more overlap at a higher level, functional enrichment analysis was performed on the rhythmically expressed genes from each dataset using the STRING database, querying KEGG pathways, Reactome pathways, and WikiPathways. Using pair-specific tested-gene backgrounds, enriched-term lists were sparse: six of the 18 pairwise annotation-category comparisons contained no enriched terms for at least one dataset, and 15 contained no more than one term in at least one dataset. Consequently, no pathway overlap remained significant after multiple-testing correction, although shared terms included circadian rhythm and several signalling-related pathways. Because this experimentally constrained background provided limited power for pathway-level comparison, we repeated the analysis using all STRING-mapped mouse genes as the background. This produced larger enriched-term sets and significant overlap in 11 of 18 comparisons after permutation testing and BH correction (Figure 3B). Both background definitions are reported in Supplementary Table 4 to show the dependence of pathway-level conclusions on the enrichment universe.

**Figure 3.**
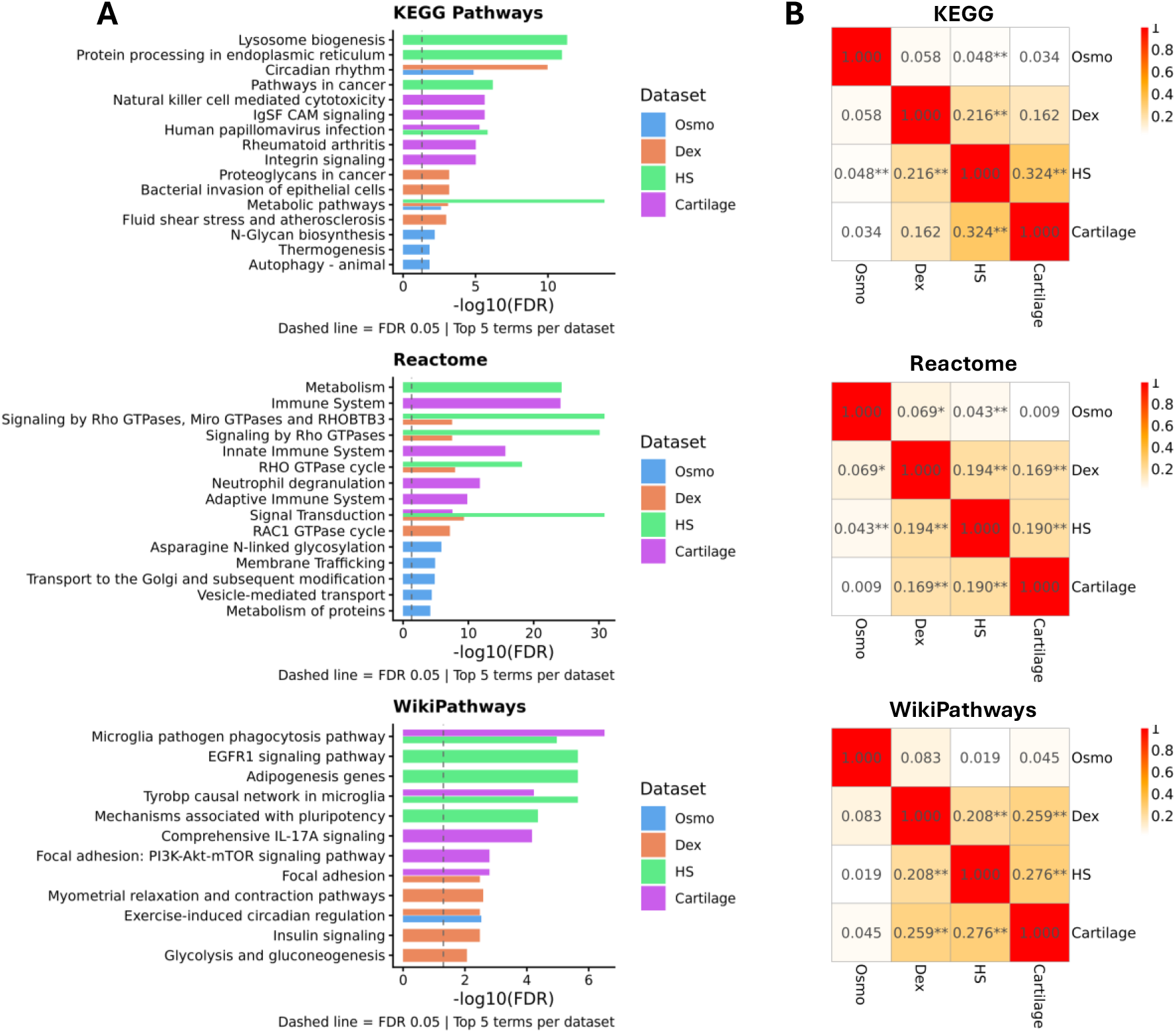
Functional enrichment overlap is limited and depends strongly on annotation category. **A**. Top 5 enrichments terms by dataset and category regardless of overlap. **B**. Jaccard index illustrating similarity of functional enrichment terms in pairwise comparisons between the chondrocyte and cartilage datasets. * BH adjusted P ≤ 0.05, ** BH adjusted P ≤ 0.01 ALT TEXT: Figures showing comparison of enriched pathways between chondrocytes treated with osmotic stress, heat shock, dexamethasone and *in vivo* cartilage.

### BodyClocks enables interactive exploration and pairwise comparison of circadian datasets

To make the analytical framework used in this study accessible across public circadian datasets, we developed BodyClocks as an interactive web resource for gene, pathway, network, and dataset-level exploration. The resource initially contains over 90 circadian transcriptomic datasets processed into a common structure, including rhythmic gene calls, peak phase estimates, functional enrichment results, and STRING protein-protein interaction networks (Figure 4A). Individual gene profiles can be visualised by direct search, table selection, enrichment term selection, or by selecting nodes within the protein-protein interaction network. Functional enrichment results, including GO Biological Process, KEGG, Reactome, and WikiPathways terms, are linked to the corresponding STRING networks, allowing pathway-associated rhythmic genes to be highlighted within their interaction context (Figure 4B and C). This allows users to assess whether genes belonging to the same biological process are connected at the protein level and whether they peak within a similar circadian time window, indicating potential phase alignment (Figure 4B and C).

**Figure 4.**
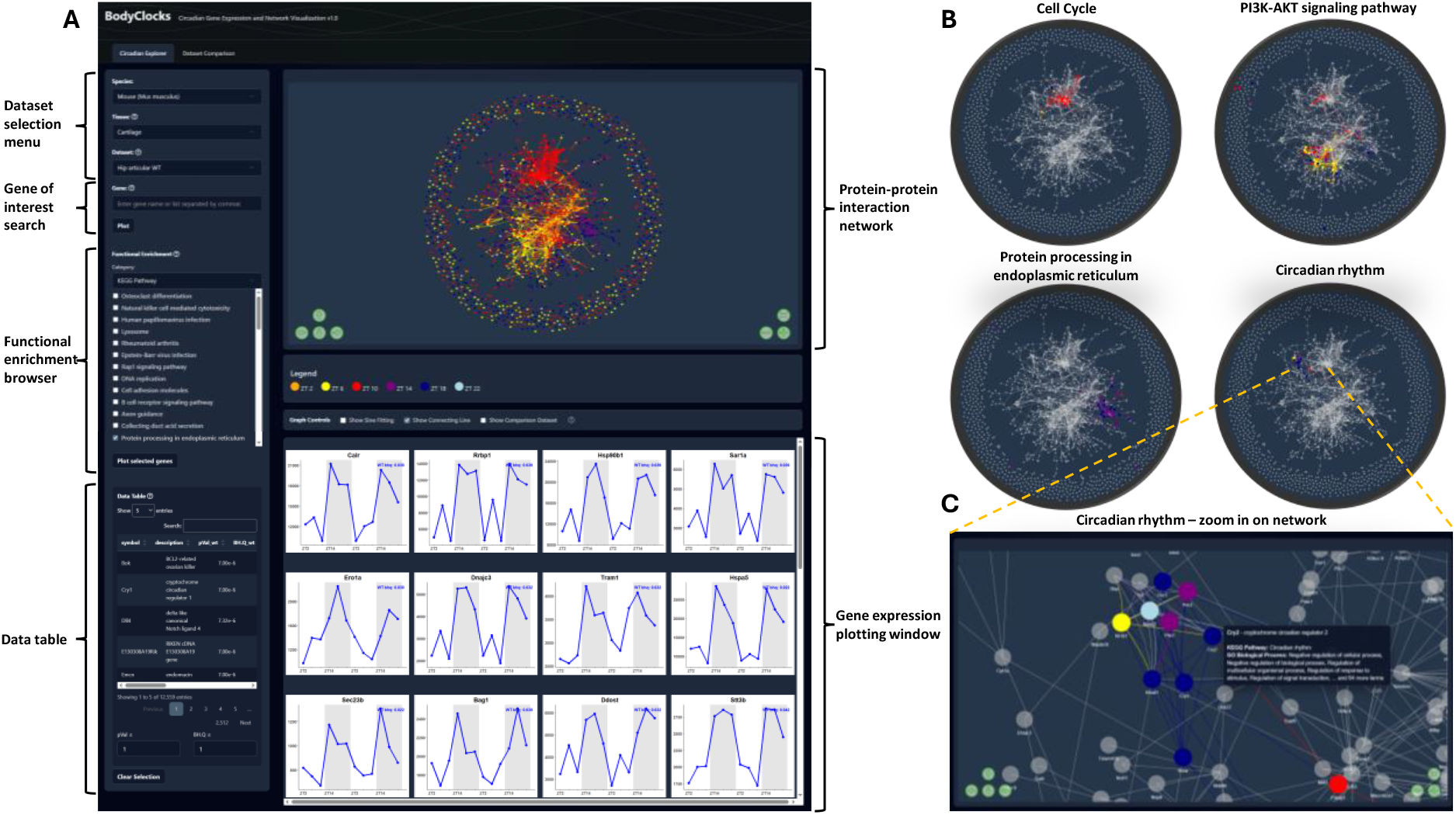
BodyClocks web resource for functional exploration of circadian transcriptomes. **A**. The Circadian Explorer module provides access to rhythmic transcriptomic datasets organised by species, tissue, and dataset. Users can search for genes, browse circadian statistics, and explore functional enrichment results across GO Biological Process, KEGG, Reactome, and WikiPathways categories. Gene expression profiles can be plotted directly from gene searches, enrichment-term selections, or the data table. **B**. Functional enrichment terms are linked to STRING protein-protein interaction networks, enabling rhythmic genes associated with a selected biological process or pathway to be highlighted in their interaction context. **C**. Interactive network exploration allows users to zoom, reposition nodes, select genes, and inspect neighbouring interactions. Selecting a node plots the corresponding gene expression profile, while hovering over a node displays gene annotation and associated enrichment terms. ALT TEXT: Graphical representation of the BodyClocks data analysis platform explaining Circadian Explorer tab features and available options including dataset selection, gene expression plotting, protein – protein association networks and functional enrichment.

Importantly, BodyClocks also implements the pairwise comparison framework used here for comparison of rhythmic transcriptomes (in our case the osmotic stress, dexamethasone, heat shock, and *in vivo* cartilage). Users can compare any two datasets from the same species to quantify rhythmic gene overlap, pathway overlap, and phase relationships among shared rhythmic genes. In this way, the chondrocyte analysis presented here serves both as a biological demonstration of zeitgeber-specific circadian output and as a worked example of the comparative workflow now available across public circadian datasets (Figure 5).

**Figure 5.**
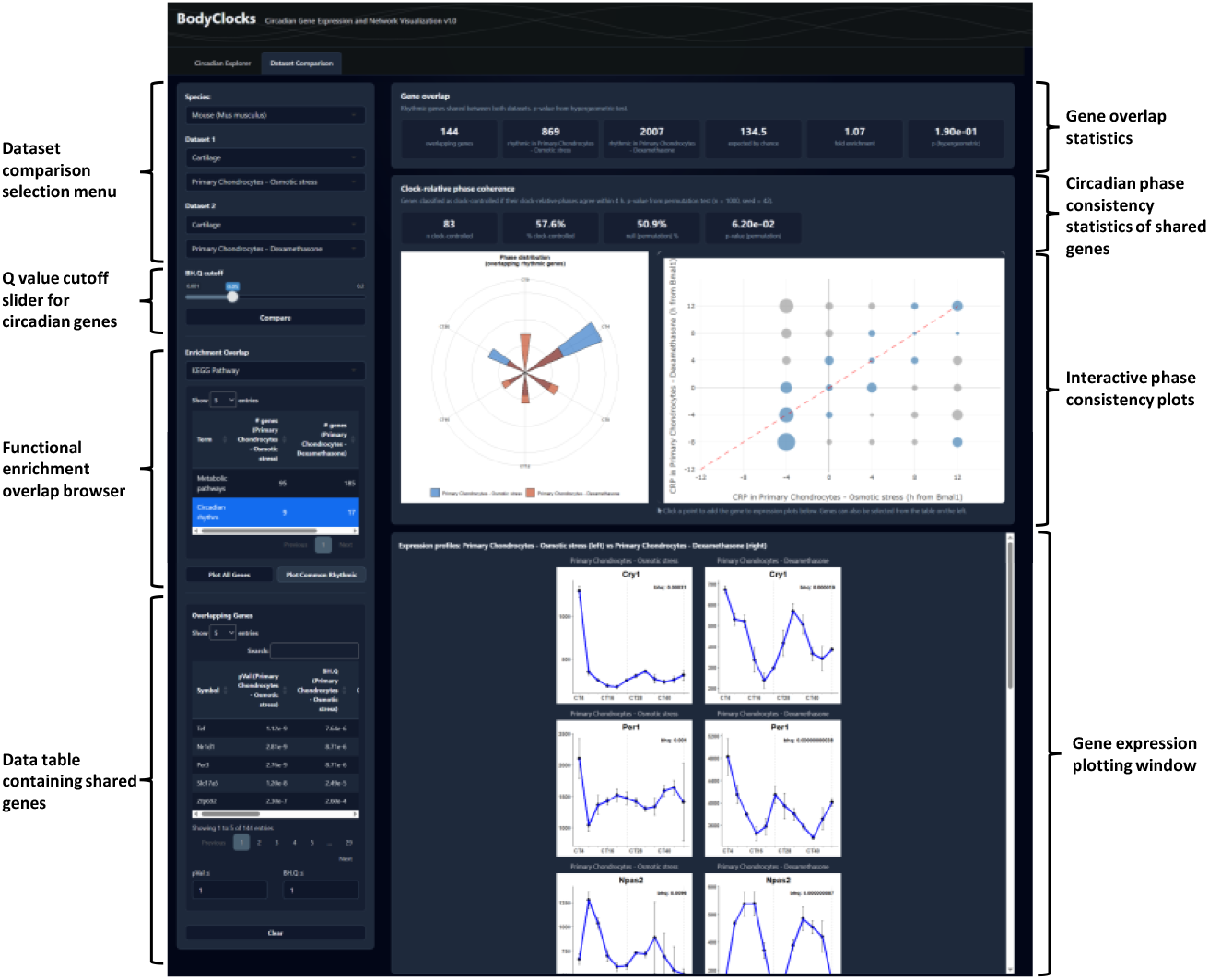
BodyClocks enables pairwise comparisons of circadian datasets. The Dataset Comparison module implements the pairwise comparison framework used in this study across BodyClocks datasets. Users can compare two datasets from the same species to quantify rhythmic gene-list overlap, assess whether the observed overlap differs from that expected by chance, and identify shared genes with similar clock-relative phase. Absolute phase relationships among shared rhythmic genes are shown in the polar plot, while clock-relative phase relationships are summarised in the bubble plot. Selecting a bubble displays expression profiles for genes at the corresponding phase intersection. The Enrichment Overlap section identifies functional terms shared between datasets and enables plotting of either shared rhythmic genes or all rhythmic genes associated with a selected term. A data table provides detailed circadian statistics for genes shared between the two datasets. ALT TEXT: Graphical representation of the BodyClocks data analysis platform explaining Dataset Comparison tab features and options including dataset selection, gene set overlap and phase coherence statistical test.

## DISCUSSION

Our study suggests that robust synchronisation of the core molecular clock is not sufficient to produce a conserved downstream circadian transcriptome that recapitulates the *in vivo* circadian programs of a given tissue. Although heat shock, dexamethasone, and osmotic stress all synchronise the chondrocyte clock, each stimulus generated a largely distinct rhythmic output with limited overlap and poor concordance with *in vivo* cartilage. The overlap between these rhythmic gene pools was minimal, performing only slightly above chance. These findings challenge the idea of a rigid, hardwired cell type-specific circadian output, suggesting instead that the peripheral clock functions as a highly flexible environmental integrator. These findings are consistent with a model in which downstream rhythmic output is shaped by interactions between the molecular clock machinery and stimulus-activated signalling pathways in cells, and the microenvironment that the cells are residing in.

The limited overlap among enriched pathways further supports the contextual nature of circadian output. Dexamethasone acts via the glucocorticoid receptor, directly driving the expression of genes possessing glucocorticoid-responsive elements, which include several clock genes such as *Per2* but also a vast number of non-clock related genes that likely interact with the clock network [36,38]. Heat shock, conversely, initiates a cellular stress response, mediated through heat shock proteins and rearrangement of the actin cytoskeleton, driving a distinct transcriptional cascade [11,39,40]. Osmotic stress, a highly relevant physiological cue for articular cartilage, results in a completely different cellular stress response involving mTOR, NFAT5, and MAP kinases [12,41–43]. Therefore, what we often define as “circadian genes” in a cell culture model may be, in reality, genes whose rhythmic expression is largely contingent upon the continuous or acute presence of a specific stimulus. This aligns with emerging literature highlighting how perturbations, such as aging [44,45], cancer [46], sleep deprivation [47] or food restriction [48–51], can fundamentally reprogram the diurnal transcriptome.

*In vivo*, chondrocytes reside in a complex, three-dimensional matrix environment where they are simultaneously exposed to diurnal temperature fluctuations, cyclic biomechanical loading, and diffusion of nutrients and systemic hormonal cues [52,53]. Because primary cells, including chondrocytes, are known to alter their transcriptome and dedifferentiate during standard *in vitro* culture [21,25], one could argue that the poor overlap between the cultured chondrocyte datasets and *in vivo* cartilage is simply an artifact of this microenvironmental shift. However, if dedifferentiation were the sole cause, we would expect a high degree of concordance among the *in vitro* models. Instead, we observed that the rhythmic gene overlap between different *in vitro* synchronisation methods was just as poor as the overlap between any *in vitro* method and *in vivo* cartilage, yielding similarly low fold-enrichment values. Indeed, the chondrocyte datasets share less rhythmic genes with each other than average *in vivo* vs *in vivo* tissue dataset comparison available in the BodyClocks platform. However, a limitation of the comparison is that the osmotic-stress and *in vivo* cartilage datasets were generated in previous studies (from our lab), and even though the same cell preparation and sampling protocols were used, may differ from the heat-shock and dexamethasone datasets. We minimised this by reprocessing datasets through a common analytical pipeline.

We have previously shown that scheduled treadmill running in mice can shift the cartilage clock nearly antiphase to the brain clock, and that the induction of osteoarthritis alters the expression of circadian clock genes [8,12]. In this context, our present results suggest that the “tissue-specific” circadian genes observed *in vivo*, similarly to the *in vitro* results, may be equally less hardwired, and more dynamically driven by the specific, complex cocktail of time cues that the tissue receives in its physiological niche, rather than a predetermined cell-specific program. Consequently, relying on a single synchronizing agent *in vitro* fails to recapitulate the physiological temporal landscape of the tissue. This suggests that the term “tissue-specific CCGs” could be more accurately defined as a “tissue niche-specific” circadian program. For example, the rhythmic transcriptome of a liver hepatocyte *in vivo* differs drastically from a joint chondrocyte not simply because their epigenetic lineages differ, but also because the liver is primarily entrained by rhythmic postprandial nutrient fluxes and humoral glucocorticoid spikes [36,48,49,51,54], whereas the articular cartilage is likely primarily entrained by the rhythmic biomechanical loading and osmotic shifts associated with the daily rest-activity cycle [12].

Given the highly contextual nature of circadian transcriptomes demonstrated here from the same cell type, our findings also highlight a limitation of interpreting circadian datasets through individual genes alone. Existing resources such as CircaDB [55] and CircaKB [56] have proven valuable for the clock community asking which gene of interest is rhythmic in which tissue, condition, or species. However, the present analysis shows that this type of gene-centric query provides only limited insight into the vast physiological information of a rhythmic transcriptome. Pairwise overlap between rhythmic gene lists can be low, and even statistically significant overlap may be only modestly greater than expected by chance. Conversely, a biologically relevant process may be rhythmically organised even when individual contributing genes differ between datasets. Therefore, understanding circadian output requires tools that ask not only whether specific genes are rhythmic, but whether rhythmic genes converge on coherent pathways, protein interaction networks, and coordinated phase relationships. BodyClocks was developed to address this need by extending circadian transcriptome exploration from single-gene lookup to pathway-, network-, and dataset-level interpretation. In the context of this study, it provides a way to examine whether zeitgeber-specific rhythmic gene lists reflect shared or divergent biological programmes. More broadly, by making the same comparative framework available across public circadian datasets, BodyClocks enables users to evaluate how rhythmic transcriptomes differ between tissues, experimental conditions, and synchronisation contexts.

While *in vitro* models remain invaluable for dissecting the design principles and molecular genetics of the core circadian clock machinery, our data indicate that single-time cue *in vitro* synchronisation models should be used with caution when inferring native tissue-level circadian outputs or prioritising downstream CCGs for physiological follow-up *in vivo*. Future chronobiology research should account for the specific clock resetting cues and local microenvironmental niche for the cell types concerned, and analytical platforms such as the BodyClocks could help disentangle the complex network of stimulus-specific circadian outputs.

## ACKNOWLEDGEMENTS

We would like to thank the Genomic Technologies Core Facility at the University of Manchester for their help with RNA sequencing, initial processing and archiving of the chondrocyte datasets. We would also like to thank Dr Steven M. Richardson, University of Manchester for critical reading of the manuscript. We would also like to thank Dr Dharshika RJ Pathiranage, University of Manchester for the continuous support of laboratory operations.

## AUTHOR CONTRIBUTIONS

Michal Dudek: Conceptualisation, Data Curation, Formal Analysis, Investigation, Methodology, Software, Supervision, Validation, Visualisation, Writing. Cátia F. Gonçalves: Investigation, Resources. Judith A. Hoyland: Conceptualisation, Funding Acquisition, Project Administration, Supervision, Writing. Qing-Jun Meng: Conceptualisation, Funding Acquisition, Project Administration, Supervision, Writing.

## DATA AVAILABILITY

The circadian time series RNA sequencing data of chondrocytes exposed to heat shock or dexamethasone is deposited at ArrayExpress under accession number E-MTAB-17163. All other data and analysis source code used in the manuscript are deposited at https://github.com/Michal0110/BodyClocks_data and Zenodo DOI: 10.5281/zenodo.21379611. BodyClocks platform source code is available at https://github.com/Michal0110/BodyClocks_app and Zenodo DOI: 10.5281/zenodo.21379114.

## REFERENCES

1. Roenneberg T, Merrow M. The Network of Time: Understanding the Molecular Circadian System. Current Biology 2003;13(5):R198–207. 10.1016/S0960-9822(03)00124-6.

2. Patton AP, Hastings MH. The suprachiasmatic nucleus. Current Biology 2018;28(15):R816–22. 10.1016/j.cub.2018.06.052.

3. Takahashi JS. Transcriptional architecture of the mammalian circadian clock. Nat Rev Genet 2017;18(3):164–79. 10.1038/nrg.2016.150.

4. Reinke H, Asher G. Crosstalk between metabolism and circadian clocks. Nat Rev Mol Cell Biol 2019;20(4):227–41.10.1038/s41580-018-0096-9.

5. Pueyo Moliner A, Ito K, Zaucke F et al. Restoring articular cartilage: insights from structure, composition and development. Nat Rev Rheumatol 2025;21(5):291–308. 10.1038/s41584-025-01236-7.

6. Okubo N, Minami Y, Fujiwara H et al. Prolonged Bioluminescence Monitoring in Mouse Ex Vivo Bone Culture Revealed Persistent Circadian Rhythms in Articular Cartilages and Growth Plates. PLOS ONE 2013;8(11):e78306. 10.1371/journal.pone.0078306.

7. Dudek M, Gossan N, Yang N et al. The chondrocyte clock gene Bmal1 controls cartilage homeostasis and integrity. J Clin Invest 2016;126(1):365–76. 10.1172/JCI82755.

8. Gossan N, Zeef L, Hensman J et al. The Circadian Clock in Murine Chondrocytes Regulates Genes Controlling Key Aspects of Cartilage Homeostasis. Arthritis & Rheumatism 2013;65(9):2334–45. 10.1002/art.38035.

9. Roenneberg T, Merrow M. The Circadian Clock and Human Health. Current Biology 2016;26(10):R432–43. 10.1016/j.cub.2016.04.011.

10. Berenbaum F, Meng QJ. The brain–joint axis in osteoarthritis: nerves, circadian clocks and beyond. Nat Rev Rheumatol 2016;12(9):508–16. 10.1038/nrrheum.2016.93.

11. Gonçalves CF, Dudek M, Pathiranage DR et al. Modulation of circadian rhythms in articular cartilage by heat pulses. Preprint, bioRxiv, 26 Jan. 2024, 2024.01.26.577368. 10.1101/2024.01.26.577368.

12. Dudek M, Pathiranage DRJ, Bano-Otalora B et al. Mechanical loading and hyperosmolarity as a daily resetting cue for skeletal circadian clocks. Nat Commun 2023;14(1):7237. 10.1038/s41467-023-42056-1.

13. Balsalobre A, Damiola F, Schibler U. A Serum Shock Induces Circadian Gene Expression in Mammalian Tissue Culture Cells. Cell 1998;93(6):929–37. 10.1016/S0092-8674(00)81199-X.

14. Chen Z, Yoo SH, Park YS et al. Identification of diverse modulators of central and peripheral circadian clocks by high-throughput chemical screening. Proceedings of the National Academy of Sciences 2012;109(1):101–6. 10.1073/pnas.1118034108.

15. Zhang EE, Liu AC, Hirota T et al. A Genome-wide RNAi Screen for Modifiers of the Circadian Clock in Human Cells. Cell 2009;139(1):199–210. 10.1016/j.cell.2009.08.031.

16. Hirota T, Lee JW, Lewis WG et al. High-throughput chemical screen identifies a novel potent modulator of cellular circadian rhythms and reveals CKIα as a clock regulatory kinase. PLoS Biol 2010;8(12):e1000559. 10.1371/journal.pbio.1000559.

17. Izumo M, Johnson CH, Yamazaki S. Circadian gene expression in mammalian fibroblasts revealed by real-time luminescence reporting: Temperature compensation and damping. Proceedings of the National Academy of Sciences 2003;100(26):16089–94. 10.1073/pnas.2536313100.

18. Balsalobre A, Marcacci L, Schibler U. Multiple signaling pathways elicit circadian gene expression in cultured Rat-1 fibroblasts. Current Biology 2000;10(20):1291–4. 10.1016/S0960-9822(00)00758-2.

19. Nagoshi E, Saini C, Bauer C et al. Circadian Gene Expression in Individual Fibroblasts: Cell-Autonomous and Self-Sustained Oscillators Pass Time to Daughter Cells. Cell 2004;119(5):693–705. 10.1016/j.cell.2004.11.015.

20. Gosselin D, Skola D, Coufal NG et al. An environment-dependent transcriptional network specifies human microglia identity. Science 2017;356(6344):eaal3222. 10.1126/science.aal3222.

21. Ma B, Leijten JCH, Wu L et al. Gene expression profiling of dedifferentiated human articular chondrocytes in monolayer culture. Osteoarthritis and Cartilage 2013;21(4):599–603. 10.1016/j.joca.2013.01.014.

22. Aguilar-Arnal L, Sassone-Corsi P. Chromatin landscape and circadian dynamics: Spatial and temporal organization of clock transcription. Proceedings of the National Academy of Sciences 2015;112(22):6863–70. 10.1073/pnas.1411264111.

23. Zhu B, Gates LA, Stashi E et al. Coactivator-Dependent Oscillation of Chromatin Accessibility Dictates Circadian Gene Amplitude via REV-ERB Loading. Molecular Cell 2015;60(5):769–83. 10.1016/j.molcel.2015.10.024.

24. Qu M, Qu H, Jia Z et al. HNF4A defines tissue-specific circadian rhythms by beaconing BMAL1::CLOCK chromatin binding and shaping the rhythmic chromatin landscape. Nat Commun 2021;12(1):6350. 10.1038/s41467-021-26567-3.

25. Caron MMJ, Emans PJ, Coolsen MME et al. Redifferentiation of dedifferentiated human articular chondrocytes: comparison of 2D and 3D cultures. Osteoarthritis and Cartilage 2012;20(10):1170–8. 10.1016/j.joca.2012.06.016.

26. Yoo SH, Yamazaki S, Lowrey PL et al. PERIOD2::LUCIFERASE real-time reporting of circadian dynamics reveals persistent circadian oscillations in mouse peripheral tissues. Proc Natl Acad Sci U S A 2004;101(15):5339–46. 10.1073/pnas.0308709101.

27. Gosset M, Berenbaum F, Thirion S et al. Primary culture and phenotyping of murine chondrocytes. Nat Protoc 2008;3(8):1253–60. 10.1038/nprot.2008.95.

28. Bolger AM, Lohse M, Usadel B. Trimmomatic: a flexible trimmer for Illumina sequence data. Bioinformatics 2014;30(15):2114–20. 10.1093/bioinformatics/btu170.

29. Dobin A, Davis CA, Schlesinger F et al. STAR: ultrafast universal RNA-seq aligner. Bioinformatics 2013;29(1):15–21. 10.1093/bioinformatics/bts635.

30. Love MI, Huber W, Anders S. Moderated estimation of fold change and dispersion for RNA-seq data with DESeq2. Genome Biol 2014;15(12):550. 10.1186/s13059-014-0550-8.

31. Zhang Y, Parmigiani G, Johnson WE. ComBat-seq: batch effect adjustment for RNA-seq count data. NAR Genom Bioinform 2020;2(3):lqaa078. 10.1093/nargab/lqaa078.

32. Thaben PF, Westermark PO. Detecting Rhythms in Time Series with RAIN. J Biol Rhythms 2014;29(6):391–400. 10.1177/0748730414553029.

33. Szklarczyk D, Kirsch R, Koutrouli M et al. The STRING database in 2023: protein–protein association networks and functional enrichment analyses for any sequenced genome of interest. Nucleic Acids Res 2023;51(D1):D638–46. 10.1093/nar/gkac1000.

34. Durinck S, Moreau Y, Kasprzyk A et al. BioMart and Bioconductor: a powerful link between biological databases and microarray data analysis. Bioinformatics 2005;21(16):3439–40. 10.1093/bioinformatics/bti525.

35. Almende BV. visNetwork: Network Visualization Using ‘vis.Js’ Library version 2.1.4. 4 Sept. 2025. https://cran.r-project.org/web/packages/visNetwork/index.html (20 May 2026, xdate last accessed).

36. So AYL, Bernal TU, Pillsbury ML et al. Glucocorticoid regulation of the circadian clock modulates glucose homeostasis. Proceedings of the National Academy of Sciences 2009;106(41):17582–7. 10.1073/pnas.0909733106.

37. Ruben MD, Wu G, Smith DF et al. A database of tissue-specific rhythmically expressed human genes has potential applications in circadian medicine. Sci Transl Med 2018;10(458):eaat8806. 10.1126/scitranslmed.aat8806.

38. Quatrini L, Ugolini S. New insights into the cell- and tissue-specificity of glucocorticoid actions. Cell Mol Immunol 2021;18(2):269–78.10.1038/s41423-020-00526-2.

39. Morimoto RI. Regulation of the heat shock transcriptional response: cross talk between a family of heat shock factors, molecular chaperones, and negative regulators. Genes Dev 1998;12(24):3788–96. 10.1101/gad.12.24.3788.

40. Gomez-Pastor R, Burchfiel ET, Thiele DJ. Regulation of heat shock transcription factors and their roles in physiology and disease. Nat Rev Mol Cell Biol 2018;19(1):4–19. 10.1038/nrm.2017.73.

41. Ortells MC, Morancho B, Drews-Elger K et al. Transcriptional regulation of gene expression during osmotic stress responses by the mammalian target of rapamycin. Nucleic Acids Res 2012;40(10):4368–84. 10.1093/nar/gks038.

42. Mordente K, Ryder L, Bekker-Jensen S. Mechanisms underlying sensing of cellular stress signals by mammalian MAP3 kinases. Molecular Cell 2024;84(1):142–55. 10.1016/j.molcel.2023.11.028.

43. Yoshitane H, Imamura K, Okubo T et al. mTOR-AKT Signaling in Cellular Clock Resetting Triggered by Osmotic Stress. Antioxidants & Redox Signaling 2022;37(10–12):631–46. 10.1089/ars.2021.0059.

44. Solanas G, Peixoto FO, Perdiguero E et al. Aged Stem Cells Reprogram Their Daily Rhythmic Functions to Adapt to Stress. Cell 2017;170(4):678-692.e20. 10.1016/j.cell.2017.07.035.

45. Sato S, Solanas G, Peixoto FO et al. Circadian Reprogramming in the Liver Identifies Metabolic Pathways of Aging. Cell 2017;170(4):664-677.e11. 10.1016/j.cell.2017.07.042.

46. Li SY, Hammarlund JA, Wu G et al. Tumor circadian clock strength influences metastatic potential and predicts patient prognosis in luminal A breast cancer. Proceedings of the National Academy of Sciences 2024;121(7):e2311854121. 10.1073/pnas.2311854121.

47. Husse J, Kiehn JT, Barclay JL et al. Tissue-Specific Dissociation of Diurnal Transcriptome Rhythms During Sleep Restriction in Mice. Sleep 2017;40(6):zsx068. 10.1093/sleep/zsx068.

48. Deota S, Lin T, Chaix A et al. Diurnal transcriptome landscape of a multi-tissue response to time-restricted feeding in mammals. Cell Metabolism 2023;35(1):150-165.e4. 10.1016/j.cmet.2022.12.006.

49. Weger BD, Gobet C, David FPA et al. Systematic analysis of differential rhythmic liver gene expression mediated by the circadian clock and feeding rhythms. Proceedings of the National Academy of Sciences 2021;118(3):e2015803118. 10.1073/pnas.2015803118.

50. Manella G, Sabath E, Aviram R et al. The liver-clock coordinates rhythmicity of peripheral tissues in response to feeding. Nat Metab 2021;3(6):829–42.10.1038/s42255-021-00395-7.

51. Vollmers C, Gill S, DiTacchio L et al. Time of feeding and the intrinsic circadian clock drive rhythms in hepatic gene expression. Proceedings of the National Academy of Sciences 2009;106(50):21453–8. 10.1073/pnas.0909591106.

52. Pueyo Moliner A, Ito K, Zaucke F et al. Restoring articular cartilage: insights from structure, composition and development. Nat Rev Rheumatol 2025;21(5):291–308. 10.1038/s41584-025-01236-7.

53. Becher C, Springer J, Feil S et al. Intra-articular temperatures of the knee in sports – An in-vivo study of jogging and alpine skiing. BMC Musculoskelet Disord 2008;9(1):46. 10.1186/1471-2474-9-46.

54. Maidstone RJ, Hunter AL, Iqbal M et al. Glucocorticoid-dependence and independence of the circadian liver transcriptome. Npj Biol Timing Sleep 2026;3(1):6. 10.1038/s44323-025-00068-8.

55. Pizarro A, Hayer K, Lahens NF et al. CircaDB: a database of mammalian circadian gene expression profiles. Nucleic Acids Res 2013;41(Database issue):D1009–13. 10.1093/nar/gks1161.

56. Zhu X, Han X, Li Z et al. CircaKB: a comprehensive knowledgebase of circadian genes across multiple species. Nucleic Acids Res 2025;53(D1):D67–78. 10.1093/nar/gkae817.

